# Inter-reader agreement of ^18^F-FDG PET/CT for the quantification of carotid artery plaque inflammation

**DOI:** 10.1101/807420

**Authors:** Kjersti Johnsrud, Therese Seierstad, David Russell, Mona-Elisabeth Revheim

## Abstract

**Background:** A significant proportion of ischemic strokes are caused by emboli from unstable atherosclerotic carotid artery plaques with inflammation being a key feature of plaque instability and stroke risk. Positron emission tomography (PET) depicting the uptake of 2-deoxy-2-(^18^F)-fluoro-D-glucose (^18^F-FDG) in carotid artery plaques is a promising technique to quantify plaque inflammation. A consensus on the methodology for plaque localization and quantification of inflammation by ^18^F-FDG PET/computed tomography (CT) in atherosclerosis has not been established. High inter-reader agreement is essential if ^18^F-FDG PET/CT is to be used as a clinical tool for the assessment of unstable plaques and stroke risk. The aim of our study was to assess the inter-reader variability of different methods for quantification of ^18^F-FDG uptake in carotid atherosclerotic plaques with a separate CT angiography (CTA) providing anatomical guidance.

**Methods and results:** Forty-three patients with carotid artery stenosis ≥70% underwent ^18^F-FDG PET/CT. Two independent readers separately delineated the plaque in all axial PET slices containing the atherosclerotic plaque and the maximum standardized uptake value (SUV_max_) from each slice was measured. Uptake values with and without background correction were calculated. Intraclass correlation coefficients were highest for uncorrected uptake values (0.97-0.98) followed by those background corrected by subtraction (0.89-0.94) and lowest for those background corrected by division (0.74-0.79). There was a significant difference between the two readers definition of plaque extension, but this did not affect the inter-reader agreement of the uptake parameters.

**Conclusions:** Quantification methods without background correction have the highest inter-reader agreement for ^18^F-FDG PET of carotid artery plaque inflammation. The use of the single highest uptake value (max SUV_max_) from the plaque will facilitate the method’s clinical utility in stroke prevention.

## Introduction

Ischemic strokes caused by thromboembolism from an unstable atherosclerotic plaque in the carotid artery can be prevented by carotid endarterectomy (CEA) [1-3]. Patients are selected for CEA based on the degree of carotid artery stenosis and presence or absence of cerebral ischemic symptoms. In recent years it has become increasingly clear that the degree of stenosis alone is not the best predictor of stroke risk. This has led to the concept of the ‘unstable plaque’ describing carotid plaques that carry high risk of stroke irrespective of the degree of artery stenosis and increased focus on factors that destabilize the plaque. Inflammation plays a key role in the development of an unstable plaque [4-6].

Positron emission tomography (PET) imaging of atherosclerosis has been rapidly evolving since the first reports of 2-deoxy-2-(^18^F)-fluoro-D-glucose (^18^F-FDG) uptake localized to the inflammatory macrophage rich areas in carotid artery plaques [7]. The goal of the imaging technique is to detect carotid plaques that are at high risk of rupture and therefore carry high risk of stroke. ^18^F-FDG PET for the detection of unstable plaques is not in clinical use [8], partly due to lack of feasible PET protocols and consensus regarding imaging procedure, method for ^18^F-FDG uptake quantification and assessment of stroke risk, although several recommendations exist [9, 10]. PET is an imaging modality with limited anatomical information, and it might therefore be challenging to define the vessel-segment-of-interest. Computed tomography angiography (CTA) is often used together with ^18^F-FDG PET when assessing patients with carotid artery stenosis, but selection of the plaque area for uptake measurements varies [11-13]. A requirement for introducing a diagnostic method into clinical routine is high inter-reader agreement. Inter-reader agreement has been studied for a few selected uptake parameters with generalized vascular inflammation [14, 15] and in patients with symptomatic carotid stenosis [12, 13], but to our knowledge no study has compared inter-reader agreement for different quantification methods.

The aim of this study was to assess inter-reader variability of different methods used for quantification of ^18^F-FDG uptake at PET/CT of carotid artery plaques.

## Materials and methods

### Study population

The study cohort consisted of forty-three patients with ultrasound-confirmed atherosclerosis with internal carotid artery stenosis ≥70% according to consensus criteria of the Society of Radiologists in Ultrasound [16]. Patient characteristics are summarized in Table 1. There were 30 men (66 ± 9 years) and 13 women (67 ± 8 years) with a mean age of 66.2 years. The study protocol conformed with the ethical guidelines of the 1975 Declaration of Helsinki and was approved by the Norwegian Regional Committee for Medical and Health Research Ethics. Written informed consent was obtained from all patients prior to study inclusion.

**Table 1.** Patient characteristics (n = 43) *.

|  |  |
| --- | --- |
| Age, years; mean ± SD | 66.2 ± 8.4 |
| Sex, male; n (%) | 30 (69.8) |
| Blood glucose, mmol·L<sup>-1</sup>; mean ± SD (range) | 6.8 ± 2.2 (4.9 - 14.9) |
| Bodyweight, kg; mean ± SD (range) | 82.4 ± 15 (55 - 110) |
| Body mass index, kg/m<sup>2</sup>; mean ± SD (range) | 27.5 ± 4.5 (19.9 - 34.8) |

\*The patient material is included in previously published studies [17, 18].

### ^18^F-FDG PET/CT examination

After a minimum of six hours fasting the patients were injected with 5 MBq/kg ^18^F-FDG and blood glucose, weight, and height were recorded. After approximately 90 minutes a two-bed position PET/CT from the base of the skull to the aortic arch was performed with 15 minutes per bed position using a hybrid PET/CT scanner (Siemens Biograph 64, Siemens Medical Systems, Erlangen, Germany). The PET images were acquired with a 256×256 matrix and the images were reconstructed to two millimetre thick slices, with four iterations/eight subsets ordered subset expectation–maximization (OSEM) algorithm and Gaussian post-reconstruction filter with 3.5 mm full width half maximum (FWHM). In addition to a non-contrast CT for attenuation correction a CTA with contrast filling of the arteries (minimum 40ml Iomeron (iodine 350mg/ml; Bracco Imaging S.P.A, Milan, Italy) or Visipaque (iodine 320mg/ml); GE Healthcare, Chicago, USA) was acquired immediately after the PET when still lying in the scanner for 16 of the 43 patients. For 24 patients CTA was performed at other radiologic departments. For three patients no CTA was available when the PET images were analysed.

### Image analyses and ^18^F-FDG quantification

The images were assessed with Hybrid Viewer 2.0 software (Hermes Medical Solutions AB, Stockholm, Sweden). Two experienced nuclear medicine senior consultants independently evaluated the ^18^F-FDG PET/CT examinations. The two readers (R1 and R2) did not undergo any joint training before assessing the images, but they agreed on how to perform the analyses. The instructions were to use the CTA as guide for drawing the region of interests (ROIs) on the fused slices (PET and non-contrast CT). The plaque was defined as vessel wall thickening and a lumen contrast-filling defect on CTA [11]. The ROIs were drawn around the entire vessel wall and lumen on all plaque-containing axial PET slices (Fig 1). For patients without CTA available, the plaque was defined as vessel wall with calcification and fat deposits in the level of the carotid bifurcation. Uptake in structures close to the plaque (e.g. lymph nodes, paravertebral muscles or salivary glands) that could falsify the plaque uptake values were excluded from the ROI. The number of plaque-containing slices for each patient was recorded. The pixel values in the PET images were converted into SUV and normalized to lean body mass [19]. SUV_max_ in all plaque containing ROIs were recorded. Background blood pool activity was obtained from four ROIs placed in the lumen of the jugular vein away from structures with ^18^F-FDG uptake but preferably in the same craniocaudal level as the plaque. The background was calculated as the mean of the SUV_mean_ in these four ROIs. Different measures of ^18^F-FDG uptake were calculated (Table 2) as previously described in detail [17]. Blood background corrected values were calculated as the ^18^F-FDG uptake values divided by the mean blood pool activity (TBR) and subtraction of the blood pool activity from the ^18^F-FDG uptake values (corrected SUV (cSUV)).

**Table 2.** Plaque ^18^F-FDG uptake measures.

| **Uptake measure** | **Description** |
| --- | --- |
| max $\text{SUV}_{\text{max}}$ | the single highest $\text{SUV}_{\text{max}}$ |
| mean $\text{SUV}_{\text{max}}$ | mean of all plaque $\text{SUV}_{\text{max}}$ |
| MDS3* | mean $\text{SUV}_{\text{max}}$ of the three contiguous slices centered on the slice with the highest $\text{SUV}_{\text{max}}$ |
| MDS5* | mean $\text{SUV}_{\text{max}}$ of the five contiguous slices centered on the slice with the highest $\text{SUV}_{\text{max}}$ |
| mean $\text{SUV}_{\text{max}}^4$ | mean $\text{SUV}_{\text{max}}$ of the four slices with highest $\text{SUV}_{\text{max}}$ |

\*MDS, most diseased segment

**Fig 1.**
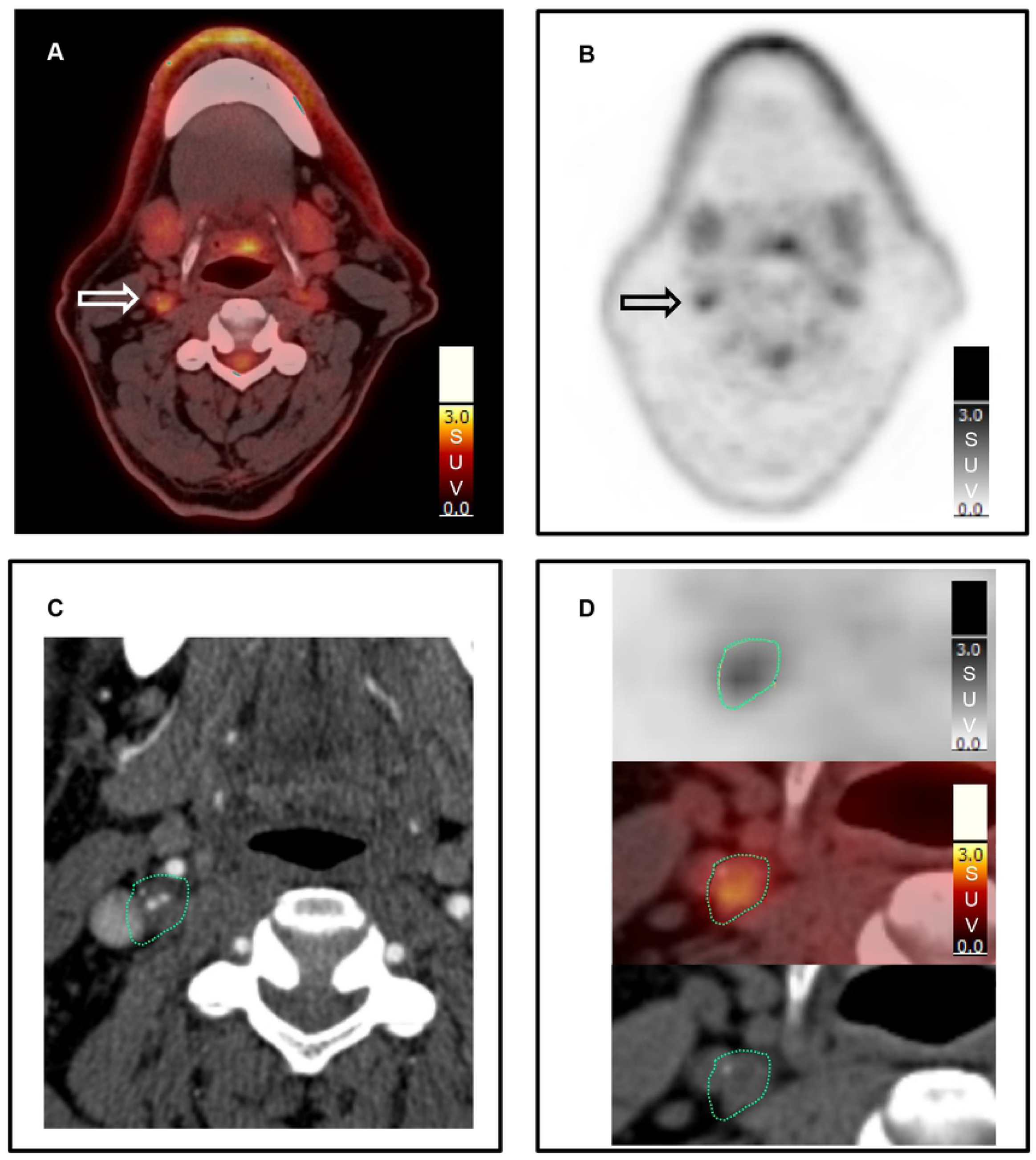
Region of interest. On each plaque-containing axial slice a region of interest (ROI) was drawn manually around the entire vessel wall including the plaque and the lumen. A (fused PET/non-contrast CT) and B (PET) show increased uptake (arrow) in the plaque in the right internal carotid artery. C shows how the plaque location on contrast enhanced CT (low attenuation plaque with thin contrast filled lumen in the centre) guides the actual drawing of the ROI (green dotted line) on the fused PET/non-contrast CT (D).

### Statistical analysis

The IBM SPSS Statistics software for Windows (version 25.0; IBM Corp., Armonk, USA) was used for data analyses. Groups of paired data were compared using the Wilcoxon signed rank test for non-normally distributed variables. Inter-reader agreement was calculated using intraclass correlation coefficients (ICC’s; model two-way random, type absolute agreement). All statistical results were considered significant when *p* < 0.05. All uptake values per patient can be found in a supplementary data file (S1 File).

## Results

The different ^18^F-FDG uptake values for the two readers are summarized in Table 3. Reader 2 identified significantly more slices as plaque containing (median; 10, range; 4-23) than reader 1 (median; 9, range; 3-18) (p = 0.001).

**Table 3.** ^18^F-FDG uptake values and intraclass correlation coefficients between the two readers (n = 43 patients)

| Quantification method | $^{18}\text{F}$-FDG uptake values | | | ICC |
| --- | --- | --- | --- | --- |
|  | Reader 1 | Reader 2 | *p* |  |
| Max $\text{SUV}_{\text{max}}$ | 1.74 (1.18 - 2.66) | 1.74 (1.20 - 2.66) | 0.304 | .979 |
| Mean $\text{SUV}_{\text{max}}$ | 1.51 (1.11 - 2.28) | 1.51 (1.06 - 2.15) | 0.687 | .973 |
| MDS3 | 1.68 (1.17 - 2.51) | 1.68 (1.19 - 2.51) | 0.400 | .978 |
| MDS5 | 1.64 (1.15 - 2.32) | 1.63 (1.17 - 2.45) | 0.438 | .972 |
| Mean $\text{SUV}_{\text{max}}^4$ | 1.68 (1.15 - 2.45) | 1.68 (1.13 - 2.45) | 0.060 | .972 |
| Background | 0.87 (0.55 - 1.26) | 0.89 (0.55 - 1.30) | 0.245 | .767 |
| TBR max $\text{SUV}_{\text{max}}$ | 1.95 (1.34 - 3.07) | 2.02 (1.34 - 2.68) | 0.314 | .792 |
| TBR mean $\text{SUV}_{\text{max}}$ | 1.72 (1.16 - 2.59) | 1.76 (1.25 - 2.37) | 0.232 | .741 |
| TBR MDS3 | 1.87 (1.26 - 2.89) | 1.97 (1.30 - 2.55) | 0.296 | .775 |
| TBR MDS5 | 1.80 (1.22 - 2.79) | 1.94 (1.24 - 2.53) | 0.241 | .769 |
| TBR mean $\text{SUV}_{\text{max}}^4$ | 1.81 (1.26 - 2.82) | 1.93 (1.31 - 2.61) | 0.358 | .758 |
| cSUV max $\text{SUV}_{\text{max}}$ | 0.83 (0.42 - 1.79) | 0.87 (0.38 - 1.67) | 0.837 | .944 |
| cSUV mean $\text{SUV}_{\text{max}}$ | 0.68 (0.20 - 1.28) | 0.68 (0.28 - 1.19) | 0.435 | .893 |
| cSUV MDS3 | 0.80 (0.33 - 1.64) | 0.79 (0.34 - 1.51) | 0.769 | .931 |
| cSUV MDS5 | 0.75 (0.28 - 1.45) | 0.76 (0.27 - 1.45) | 0.595 | .916 |
| cSUV mean $\text{SUV}_{\text{max}}^4$ | 0.74 (0.32 - 1.58) | 0.77 (0.35 - 1.45) | 0.975 | .919 |

Data are given as median (range). *P*-value from Wilcoxon signed ranks test. SUV, standardized uptake value; MDS, most diseased segment; TBR, target-to-background ratio; cSUV, background subtracted SUV; ICC, intraclass correlation coefficient.

There were no differences in ^18^F-FDG uptake between the two readers (Table 3). The ICC for the different ^18^F-FDG quantification methods was highest for uncorrected SUVs (0.97-0.98) followed by cSUVs (0.89-0.94) and TBRs (0.74-0.79), and 0.77 for the background blood pool (Table 3). The differences in the median for the uptake values between the readers ranged from 0.00 and 0.01 for the uncorrected SUVs to 0.04-0.14 for TBRs (0.14 for TBR MDS5). The difference for the background value was 0.02 (Table 3).

Fig 2 shows the differences in max SUV_max_ and mean SUV_max_ for individual patients for the two readers without background correction (A and B), and the corresponding values when the ^18^F-FDG uptake is corrected for background blood pool by division (TBR; 2C and 2D) and by subtraction (cSUV; 2E and 2F). The difference in venous background is shown in Fig 2G. The difference between the readers is highest for the uptake values corrected for background blood pool by division (2C and 2D). and lowest for the uptake values without background correction (2A and 2B).

**Fig 2.**
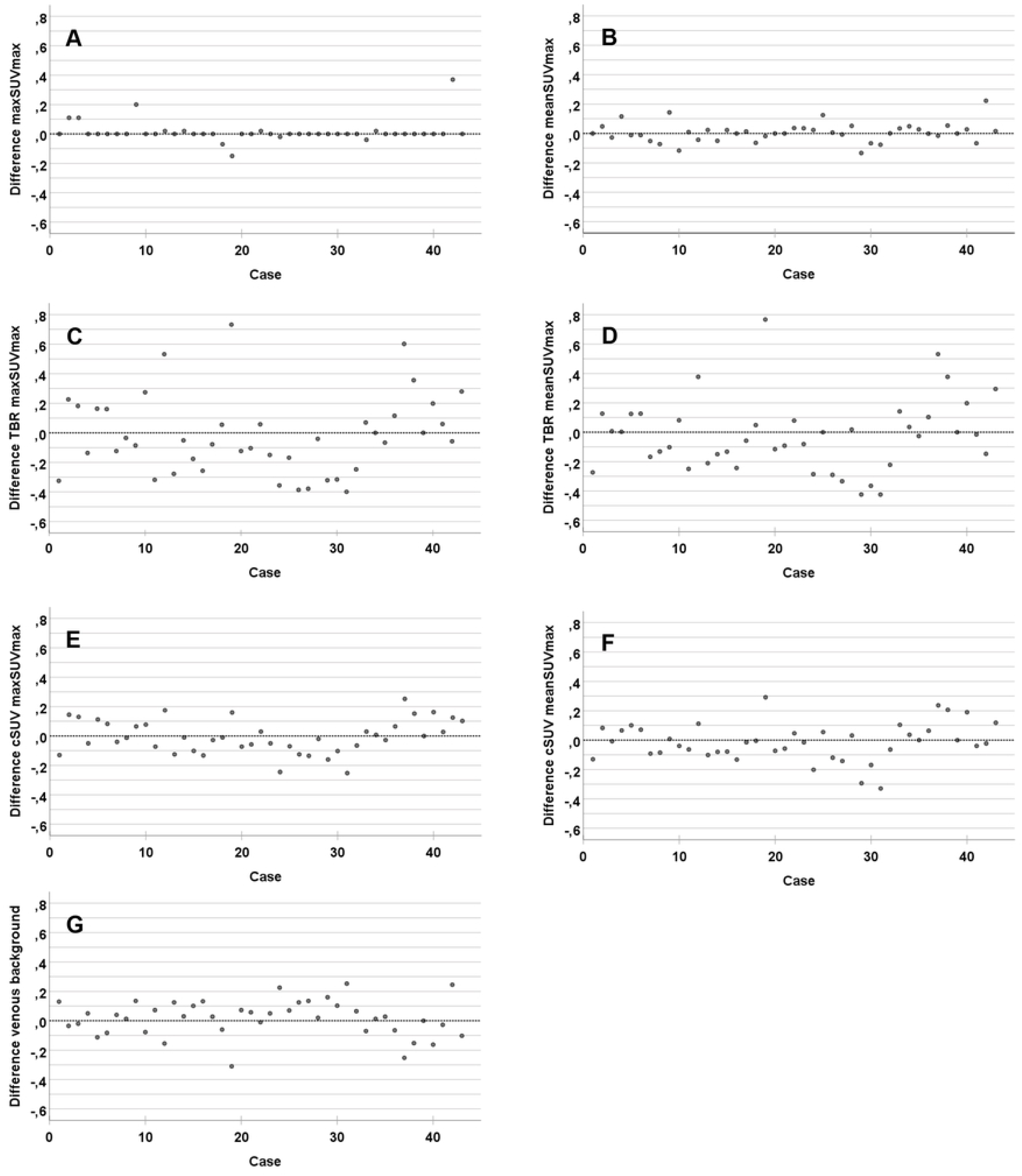
Inter-reader difference for the ^18^F-FDG quantification methods. Difference between the readers (R2 minus R1, y-axis)) for the included patients (x-axis). Max SUV_max_ (A), mean SUV_max_ (B), TBR max SUV_max_ (C), TBR mean SUV_max_ (D), cSUV max SUV_max_ (E), cSUV mean SUV_max_ (F), and venous background (G).

## Discussion

In this study we found high inter-reader agreement between different methods for ^18^F-FDG uptake quantification of inflammation in high grade carotid artery stenosis. The inter-reader agreement was highest for the methods without background correction. Two studies in patients with carotid stenosis supports our finding that methods without correction for background blood activity have higher inter-reader agreement than background corrected values: Kwee et al. [12] reported an ICC of 0.61 for TBR mean SUV_max_ and 0.65 for TBR max SUV_max,_ and Marnane et al. [13] found an ICC of 0.99 for mean SUV_max_.

In our study the highest ICC was found for max SUV_max_ (0.98). For the methods without background blood pool correction only 12% of the max SUV_max_ and 14% of the mean SUV_max_ measurements differed with more than ± 0.10 (Fig 2A, B). Patient number 42 is an outlier with an inter-reader difference of 0.38. This is probably due to different delineations of the plaque ROIs as this patient had high uptake in neighbouring muscle (Fig 3). Reader 1 can have excluded more of the plaque ROIs to be sure to avoid spill-in activity than reader 2. The problem with spill-in from neighbouring structures is due to the relatively low spatial resolution of PET combined with unspecific uptake of ^18^F-FDG.

**Fig 3.**
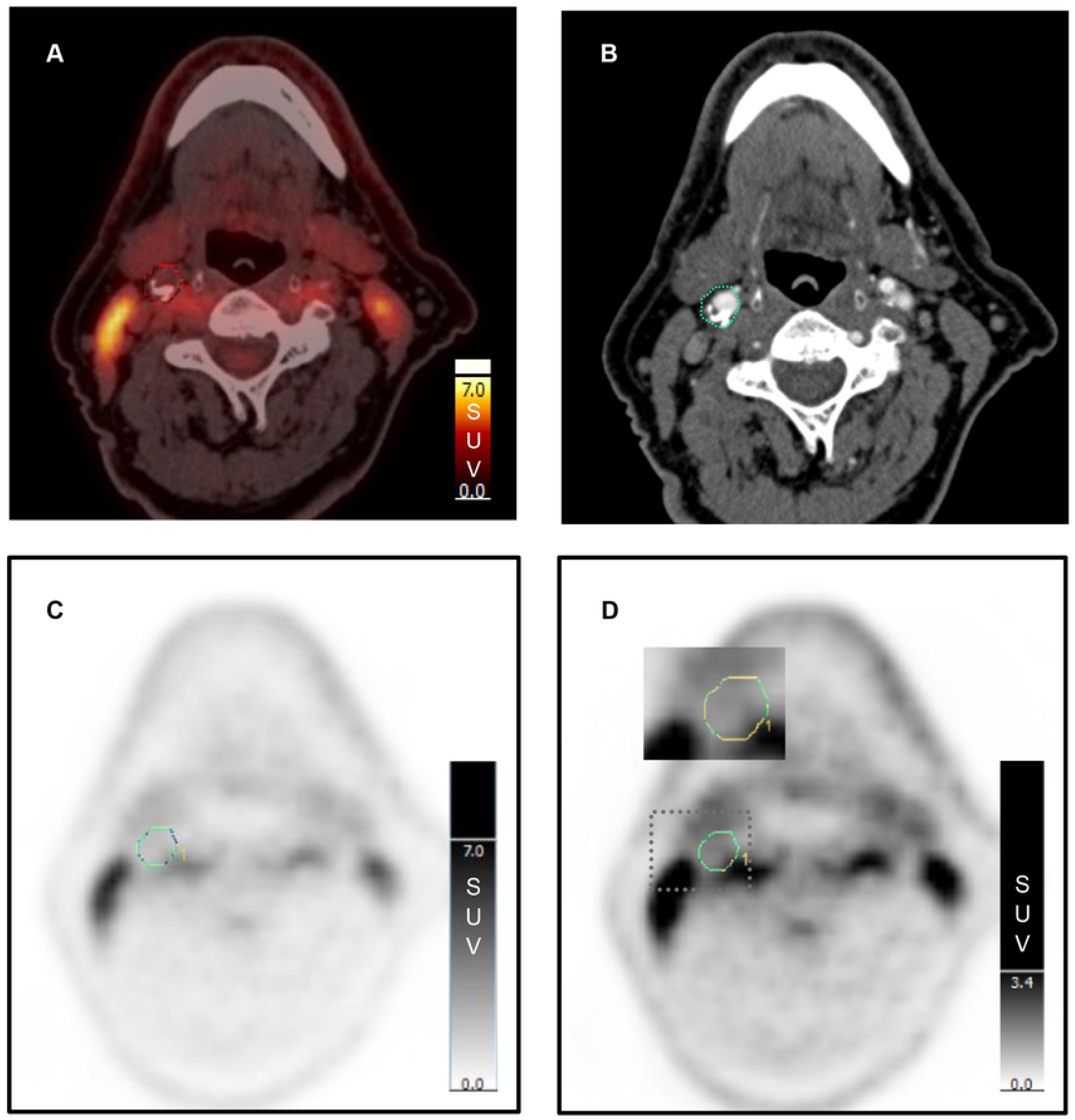
Spill-in activity. Fused image of non-contrast CT and PET (A) and contrast enhanced CT (B) show a plaque in the level of the right carotid bifurcation with low uptake but with high uptake in nearby muscles. PET with normal intensity on the SUV scale (C) and PET with high intensity on the SUV scale (D) show that ^18^F-FDG uptake from nearby muscle activity influences the ROI around the plaque (inserted picture at 4 to 5 o’clock position).

For the background corrected values, the difference was larger with 40% of TBR max SUV_max_ and 30% of TBR mean SUV_max_ having a difference of ± 0.25 or more (Fig 2C, D). In our previous study exploring ^18^F-FDG-uptake in symptomatic versus asymptomatic patients [18] the difference in median mean SUV_max_ between the groups was 0.32 (1.75 versus 1.43). In two studies using TBR max SUV_max_ as uptake parameter the difference was found to be 0.19 and 0.29 [20, 21]. Thus, methods with reader difference of 0.25 prohibit differentiation between symptomatic and asymptomatic patients.

We found an ICC for background blood pool activity of 0.77. This discordant assessment of background blood pool activity introduces variation in TBR and cSUVs due to methodology rather than biology. The background blood pool activity in our study was obtained from four ROIs within the lumen of the jugular vein preferably in the same craniocaudal level as the plaque. The vena jugularis has a small diameter and it was often challenging to draw reproducible ROIs within the vein that excluded contribution from neighbouring structures. In a ^18^F-FDG PET study of generalized vascular inflammation in which the background blood pool activity was obtained from eight ROIs in the jugular vein the ICC for TBR mean SUV_max_ of the carotid arteries was 0.94-0.96 [14]. This suggests that including data from more slices or from a larger vessel segment such as the vena cava superior or atria of the heart could have reduced the inter-reader variability of measuring the blood pool activity. In this study the two readers also had trained together by co-reading several pilot studies before they established an analysis protocol [14]. This is optimal for research studies, but hard to accomplish in larger trials where the readers often are located in different departments.

There is a large amount of studies that quantifies the ^18^F-FDG uptake in the vessel wall of patients with suspected generalized vascular inflammation (atherosclerosis not necessarily confirmed by other imaging methods). Although our findings cannot automatically be generalized, one might question the need for background correction for these patients.

Reader 2 included significantly more plaque-containing slices than reader 1. This did not reduce the ICC of the ^18^F-FDG measurements, supporting that the plaque slices with the highest uptake values all were included in both readers plaque area and that the number of slices included in the plaque area has minimal influence on mean SUV_max_. Our interpretation of this finding is that the plaque inflammation we can detect with ^18^F-FDG PET is homogeneously spread out, and also present in the extreme tails of the plaque. This was also one of our main findings when we explored associations between different ^18^F-FDG uptake parameters and plaque inflammation at histopathology [17]. Furthermore, this is in accordance with the study results from Kwee et al. [12] who found a strong correlation between TBRs of ipsilateral symptomatic plaques and contralateral asymptomatic plaques and supports the hypothesis that plaque inflammation is systemic to some extent.

A strength of our study is a relatively large patient population with a wide range of uptake values (max SUV_max_ from 1.18 to 2.66) representing low to high plaque inflammatory activity confirmed by histology [17].

In conclusion, our study confirms the reproducibility of quantification of ^18^F-FDG uptake in carotid artery plaques and supports the superiority of quantification methods that do not include blood pool background. The ICC was highest for max SUV_max_ (the single highest uptake value within the plaque) and thus, our suggestion is to further explore this parameter for atherosclerosis imaging.

## Supporting information

**S1 File**. SPSS file with individual ^18^F-FDG uptake measurements.

## Author Contributions

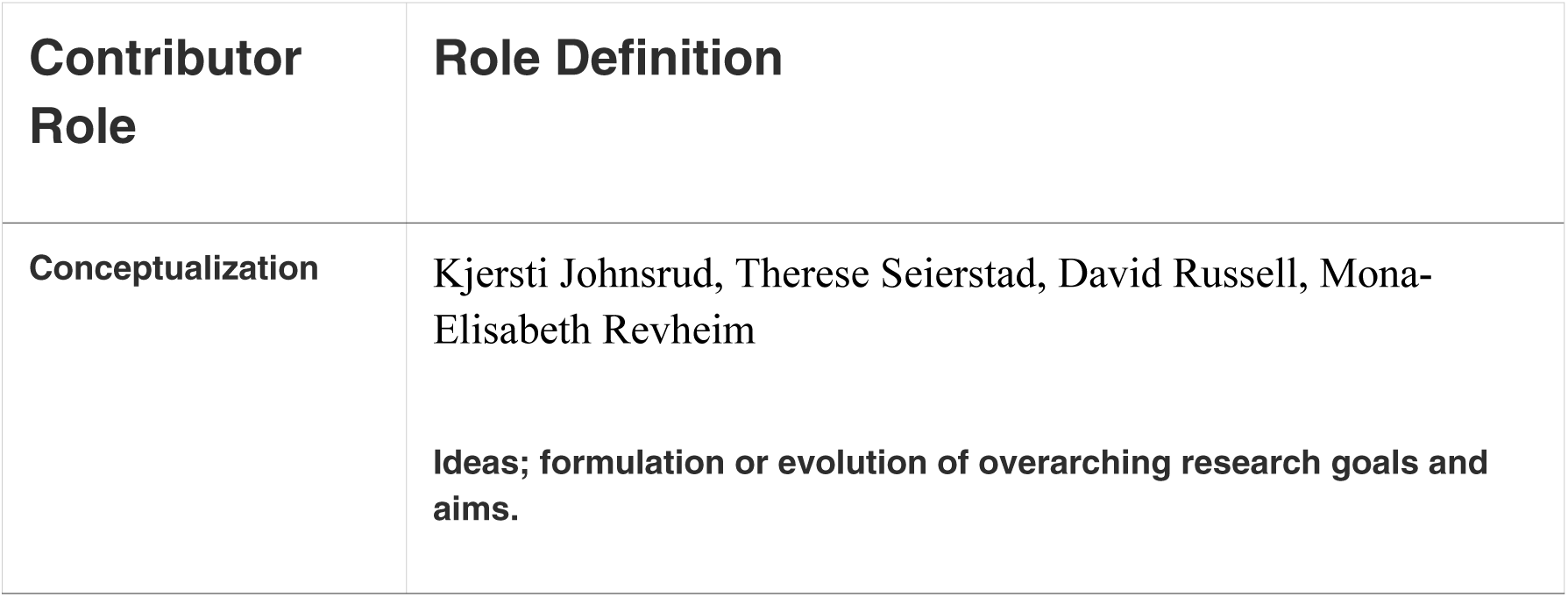
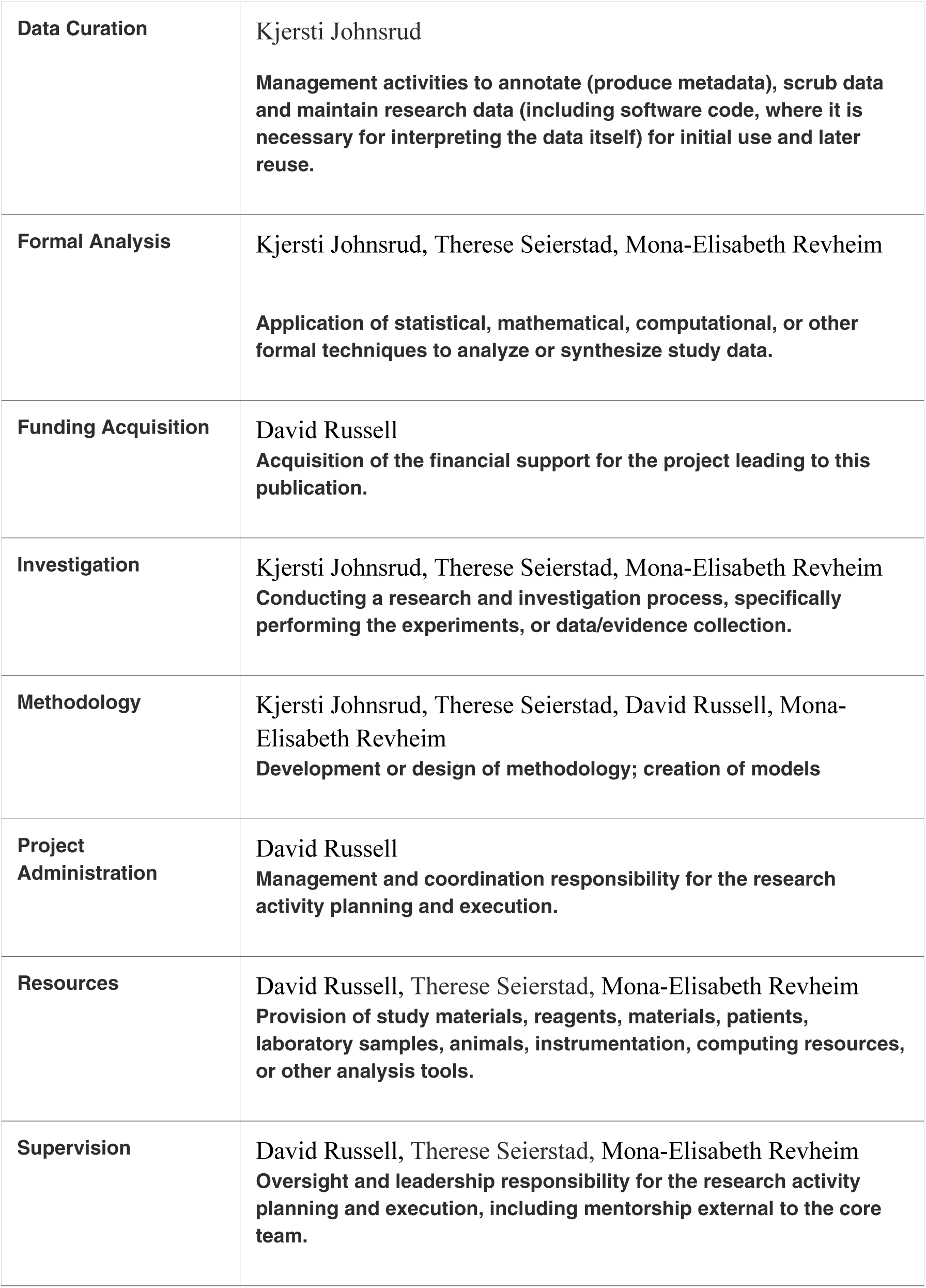
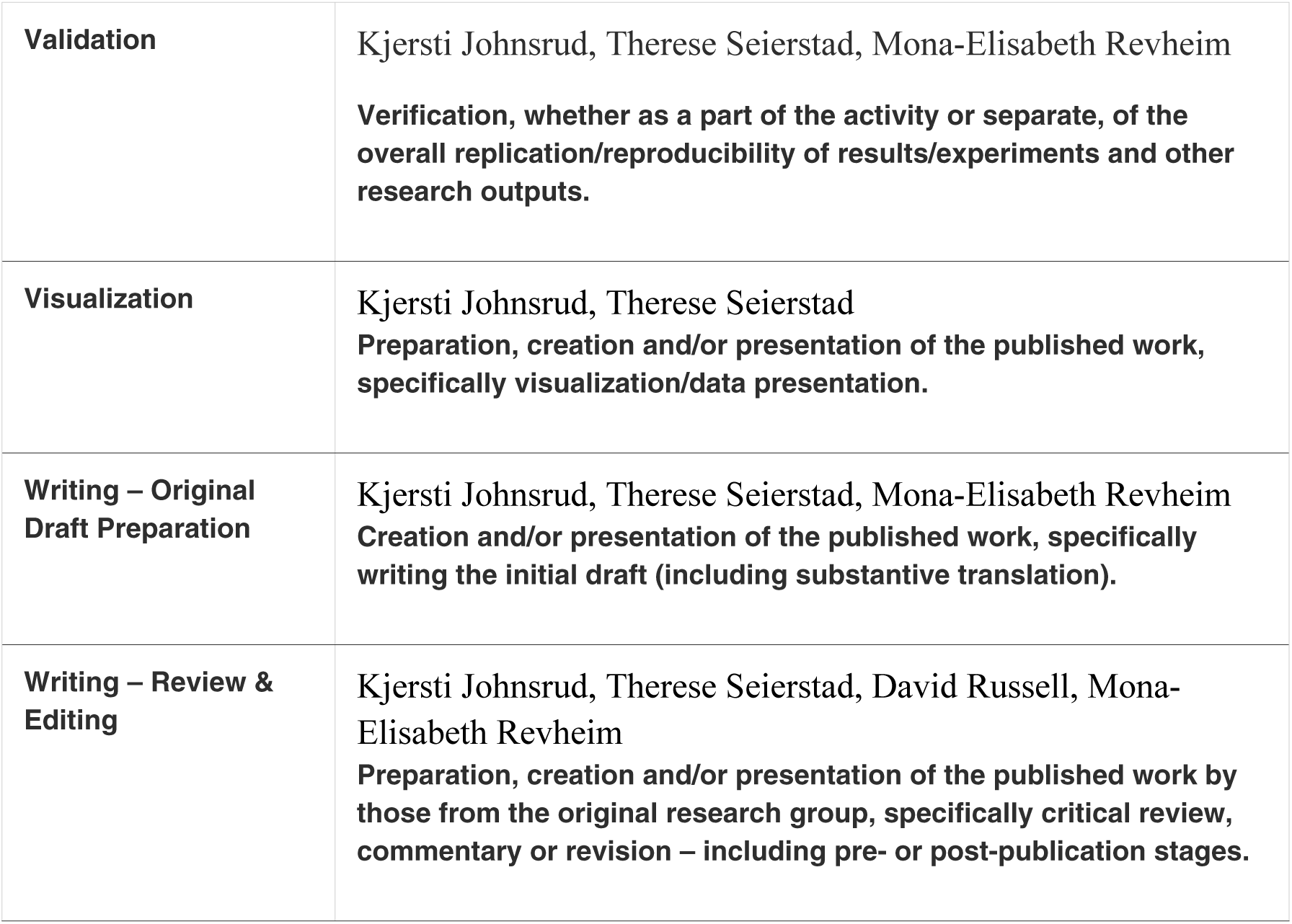

